# Predicting hERG repolarization power at 37°C from recordings at room temperature

**DOI:** 10.1101/2023.04.13.536707

**Authors:** Barbara B.R. Oliveira-Mendes, Malak Alameh, Jérôme Montnach, Béatrice Ollivier, Solène Gibaud, Sylvain Feliciangeli, Florian Lesage, Flavien Charpentier, Gildas Loussouarn, Michel De Waard, Isabelle Baró

**Affiliations:** Nantes Université, CNRS, INSERM, l’institut du thorax, F-44000 Nantes, France; Labex ICST, Université Côte d’Azur, INSERM, Centre National de la Recherche Scientifique, Institut de Pharmacologie Moléculaire et Cellulaire, Valbonne, France

**Keywords:** repolarization power, diagnostic testing, genetic variant, hERG ion channel, temperature, QT syndromes, translational medicine

## Abstract

Mutations in the *KCNH2* gene cause long or short QT syndromes (LQTS or SQTS) predisposing to life-threatening arrhythmias. *KCNH2* encodes for the voltage-gated K^+^ channel hERG involved in the late repolarization phase of the cardiac action potential (AP). For the last decades, sequencing *KCNH2* has provided a plethora of variants associated or not with clear pathological cardiac phenotypes. Identifying pathogenic or likely pathogenic variants from the benign ones would provide useful information to clarify the genetic background of LQTS patients and relatives, and to stratify the risk of adverse events. In face of a wide spectrum of hERG biophysical defects, we looked for a way to summarize the net loss or gain of function in a unique index. In a previous work, we defined as the repolarization power the time integral of the K^+^ currents developed during an AP clamp. Here, with the aim of accelerating the functional characterization of hERG variants using automated patch-clamp, we adapted the AP-clamp protocol to establish, at room temperature, at which the recording success rate is high, a repolarization power index, as reliable and informative as the one measured at physiological temperature. We also illustrate that the repolarization power determined at room temperature is predictive of the repolarization power at physiological temperature for 2 pathogenic hERG variants with different biophysical dysfunctions.

## Introduction

The duration of the QT interval on the surface electrocardiogram (ECG) is a representation of the repolarization time in cardiac ventricles. QT intervals vary as a function of various physiological (age, heart rate, hormones…) or pathophysiological (heart disease, fever, or drug intake) factors^1^. Normal QT intervals, when corrected by heart rate (QTc), are below 470-480 ms. When these QTc intervals are markedly prolonged, generally to values greater than 550–600 ms, polymorphic ventricular tachycardia such as torsade de pointes, elicited by emotions or physical activity, may occur and can degenerate to fatal arrhythmias such as ventricular fibrillation. Excessive QTc prolongations may have various origins: mutations in genes related to ion channels, as in the congenital long QT syndrome (cLQTS)^2^, or upon exposure to stressing environmental conditions as in the acquired long QT syndrome (aLQTS)^3^. aLQTS is frequently provoked by the intake of QT-prolonging drugs, many of them of common use^4^. In this case, reversion back to normal generally follows withdrawal of the causative trigger.

Three genes encoding ion channel pore-forming subunits have been identified as responsible for up to 75% of the cLQTS cases: *KCNQ1* (K_V_7.1 channel, 30–35% of LQTS cases), *KCNH2* (hERG *alias* K_V_11.1 channel, 25–30%) and *SCN5A* (Na_V_1.5 channel, 5–10%)^5–7^. A third of aLQTS patients carry cLQTS mutations, those on *KCNH2* being more common^8^. Physiologically, the two voltage-gated K^+^ channels K_V_7.1 and hERG, are responsible, as pore-forming channel α-subunits, for delayed outward currents, I_Ks_ and I_Kr_, respectively, involved in the cardiac action potential (AP) repolarization^9^. Their regulation contributes to AP duration adaptation to heart rate. A reduction in outward current and/or an increase in inward current, a condition called reduced repolarization reserve^10^, leads to AP duration lengthening, underlying the QT interval prolongation. About a thousand of cLQTS-associated *KCNH2* missense variants have been reported^11^.

Finally, gain-of-function hERG mutants have been detected in patients with shorter QT duration (≤ 360 ms) on the ECG, presenting symptoms varying from atrial to ventricular fibrillation and sudden death^12, 13^.

For the last decades, sequencing *KCNH2* has provided a plethora of variants associated or not with clear pathological cardiac phenotypes^14, 15^. Distinguishing pathogenic or likely pathogenic variants from the benign ones is a critical information to clarify the genetic background of LQTS patients as well as their relatives. Today, up to 70% of *KCNH2* detected missense variants are still classified as variants of unknown significance (VUS) or with conflicting interpretations^16^ according to the American College of Medical Genetics and the Association for Molecular Pathology classification guidelines^17^. Various databases gathering data from LQTS patients have been continuously developed, as Bamacoeur in France^18^, but they all have limited information regarding the functional impact of the variants at the molecular level. In addition, when functionally investigated, the heterogeneity of the models and approaches prevents from clearly classifying them as illustrated by different studies focusing on the same variants ^19–24^. Therefore, in an attempt to standardize the assessment of hERG variants molecular pathogenicity, we have recently designed a hERG phenotyping pipeline^24^. In order to speed up the molecular phenotyping, we designed a new voltage-clamp protocol to determine more than 10 biophysical parameters of hERG current (I_hERG_) in 35 seconds. To easily summarize the net functional effect of the modified biophysical parameters, we defined a new, simple and unique index, we called repolarization power, to grade the molecular pathogenicity of hERG channels. This index is the time integral of the current density recorded during an AP-shaped voltage-clamp stimulation (AP-clamp), therefore proportional to the total amount of K^+^ ions crossing the membrane during each AP.

The development of the automated patch-clamp allows now to studying up to 384 cells in parallel, boosting the variant functional investigation speed. However, the aforementioned repolarization power, representative of *in vivo* hERG contribution to repolarization, has to be established at physiological temperature. Indeed, the K^+^ current conducted by hERG channels (I_hERG_) depends on temperature but in a complex manner ^25, 26^. Recently, Lei and collaborators have developed a short and informatively rich protocol to extensively study hERG behavior and its temperature dependence^27, 28^. Namely, they associated a simple Hodgkin-Huxley kinetic model, an Eyring formulation of the temperature dependence in the model, and a 15-s stimulation protocol (staircase protocol) designed for any patch-clamp set-up, including high-throughput automated systems. All these studies showed that the relative occupancy of the channel open state (total K^+^ conductance) and the rates of activation, deactivation, inactivation, and recovery from inactivation have different temperature sensitivities, the activation being far more temperature sensitive than inactivation^25–28^. Furthermore, Lei also showed that experimental estimations of temperature coefficients (Q_10_) are highly protocol dependent. Therefore, estimation of the effects of hERG variants on repolarization *in vivo* requires to ideally work at 35-37°C.

On the other hand, as reported by Rajan and colleagues, in CHO, CV1, or HEK cells studied with the Nanion NPC-16 Patchliner Quattro, the patch-clamp success rate, defined in this study as achieving and maintaining a seal resistance above 200 MΩ, at 35°C could be as low as ∼15% compared to ∼80% at 25 and 15°C^29^. Thus, high temperature represents a limit to obtain high success rates in patch-clamp experiments and will therefore preclude high-throughput evaluation of all hERG variants.

In this report, our aim was to optimize the functional characterization of hERG variants by circumventing these two opposite constraints (physiological temperature and seal resistance in the GΩ range). To do so, we adapted the AP-clamp protocol^24^ in such a way that it could be used at room temperature to predict the reference repolarization power of hERG channels at 37°C.

Since hERG gating kinetics are slower at lower temperature, we attempted to mimic the hERG current profile observed at 37°C, by applying slower variation of voltage than the one used for AP-clamp at 37°C. Using AP-shaped voltage stimulations of various time scales applied at different temperatures, we determined that a unique temperature coefficient of 2 can be used to generate a current profile at room temperature, that matches the one generated at physiological temperatures. We also illustrate that the repolarization power determined at room temperature is predictive of the repolarization power at physiological temperature for two pathogenic hERG variants with different biophysical dysfunctions.

## Material and methods

### Cell culture

HEK293 cells were cultured in Dulbecco’s modified Eagle’s medium (Gibco, France) supplemented with 10% fetal calf serum (Eurobio, France), 4.5 g/L glucose, 2 mmol/L L-glutamine, 100 U/mL penicillin, 100 µg/mL and streptomycin (Corning, France) at 5% CO_2_, maintained at 37°C in a humidified incubator. HEK293 cells stably expressing the human hERG channel (Bioprojet, France) were cultured in the same medium with 400 µg/mL G418 (Thermo Fisher Scientific, France), at 5% CO_2_, maintained at 37°C in a humidified incubator. The cell lines were confirmed to be mycoplasma-free (MycoAlert, Lonza, France).

### hERG plasmid transfection

In a first attempt of mutant hERG current recordings using the automated patch-clamp system, HEK293 cells were transfected by electroporation using the MaxCyte STx system (MaxCyte Inc., MD, USA), using 30 µg plasmid per 100 µl Hyclone buffer^30^. For conventional patch-clamp experiments, the FuGENE 6 transfection reagent (Promega, WI, USA) was used to transfect wild-type (WT, protein sequence: NP_000229), p.R328C and p.D591H hERG plasmids. HEK293 cells (passages 24-30) were cotransfected, using 6 µL of FuGENE 6, with 1.6 µg WT or mutant pcDNA5/FRT/TO Opti-hERG and 0.4 µg peGFP (Clontech, France) for fluorescence-based cell selection in microscopy^24^. Twenty-four hours after transfection, cells were trypsinized, diluted and plated on 35-mm dishes to obtain isolated cells for patch-clamp experiments.

### High-throughput automated electrophysiology

Temperature effects on hERG channels were investigated on HEK293 cells stably expressing the hERG channel using the automated patch-clamp system SyncroPatch 384PE (Nanion Technologies, Germany). Cells were detached with accutase (Innovative Cell Technologies, Inc., CA, USA) and floating single cells were diluted (∼300,000 cells/mL) in medium containing (in mmol/L): NaCl 140, KCl 4.0, CaCl_2_ 2.0, MgCl_2_ 2.0, glucose 5.0 and HEPES 10 (pH 7.4, osmolarity 290 mOsm). Prior to recordings, dissociated cells were gently shaken at 200 rpm in a cell hotel reservoir at 10°C. NPC-384T 1x L-Type chips with typical single-hole and resistance of 2.4 ± 0.02 MΩ (n = 384) were used for recordings. Voltage stimulation and whole-cell recording were achieved using the PatchControl384 v1.5.3 software (Nanion Technologies) and the Biomek v1.0 interface (Beckman Coulter, France). The intracellular solution contained (in mmol/L): KCl 10, KF 110, NaCl 10, EGTA 10 and HEPES 10 (pH 7.2, osmolarity 280 mOsm), and, after seal formation, the extracellular solution contained (in mmol/L): NaCl 140, KCl 4.0, CaCl_2_ 2.48, MgCl_2_ 1.69, glucose 5 and HEPES 10 (pH 7.4, osmolarity 298 mOsm). After cell catch, seal, whole-cell formation and compensation, liquid application, and data acquisition were all performed sequentially and automatically. Whole-cell experiments were done at temperatures ranging from 22°C to 37°C. An AP stimulation protocol made of 6 voltage steps and ramps was designed to mimic the human ventricular AP from the O’Hara and Rudy model^31^ of sub-epicardial ventricular action potential at 37°C (Supplemental figure 1). Automated patch-clamp imposes the use of intracellular fluoride and high extracellular Ca^2+^ concentration (10 mmol/L) to obtain seals between the cell and the hole bored at the glass bottom of the recording well, close to 1 GΩ (Gigaseal). Extracellular Ca^2+^ concentration, after dilution, was kept relatively high (2.48 mmol/L) to maintain acceptable seal quality. To correct the screening effects of external Ca^2+^ on the surface charge of the membrane^32, 33^, and the uncompensated liquid junction potential^34^, the stimulation potential was shifted by +32 mV. On the other hand, the holding potential was set to −80 mV, to allow full recovery from inactivation of hERG current (see Supplemental figure 2 for intermediary optimization steps and Figure 2A, the dashed line representing the optimized AP). AP stimulation was repeated 4 times at a frequency of 1 Hz to ensure stability of the parameters.

**Supplemental figure 1:**
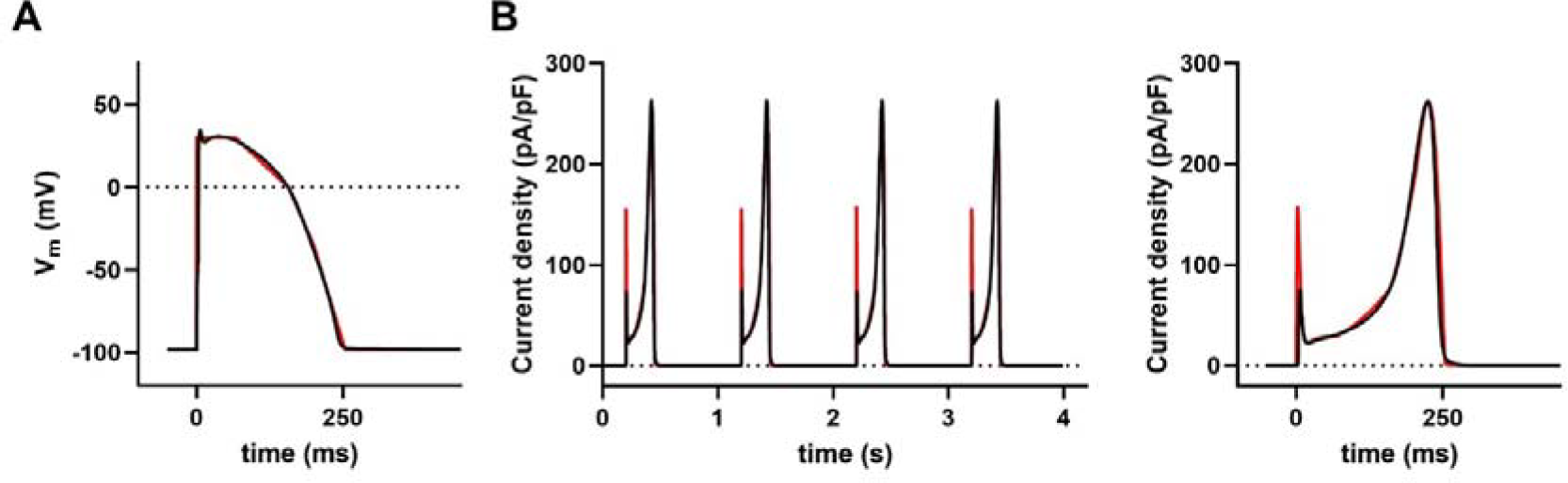
The I_Kr_ current (B, red), generated *in silico*^35^ using the simplified AP (A, red) as stimulation, adequately overlaps with the one (B, black) using the AP generated by O’Hara and Rudy model (A, black)^31^.

**Supplemental figure 2:**
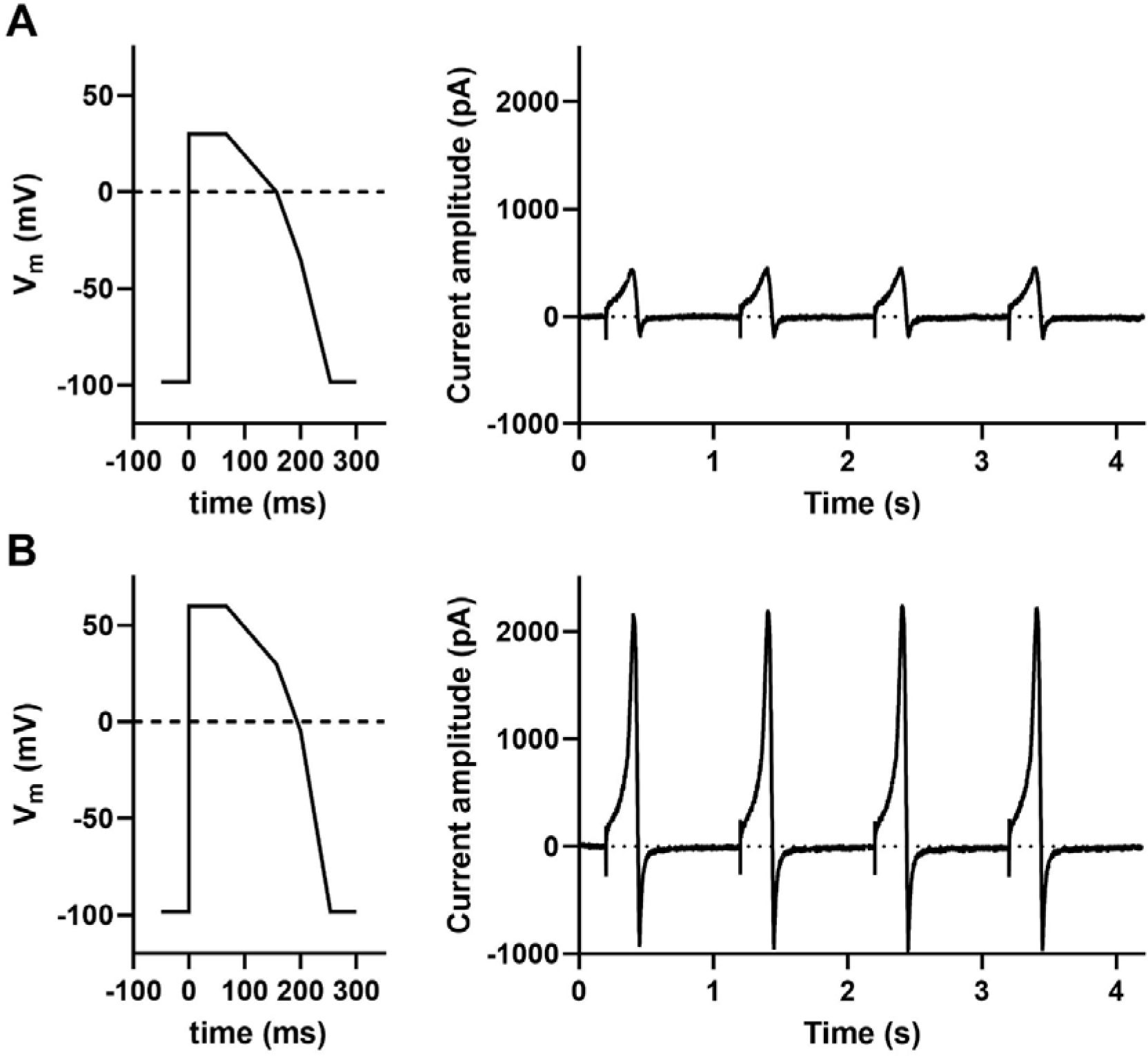
Optimization of the AP stimulation protocol for automated patch clamp. (A) Original AP (made of 6 voltage steps and ramps) mimicking the human sub-epicardial ventricular AP from the O’Hara and Rudy model^31^. (B) After correction of the screening effects of external Ca^2+^ on the surface charge of the membrane and compensation of the liquid junction potential, the higher overshoot allowed development of a much larger K^+^ current (same cell as A). In this example, the holding potential (HP) was still set at −100 mV and a transient inward current due to hERG deactivation was recorded at the end of the AP repolarization. Therefore, HP of the final AP stimulation protocol was set to −80 mV, limiting the inward K^+^ current but still allowing full recovery of hERG current.

**Figure 2:**
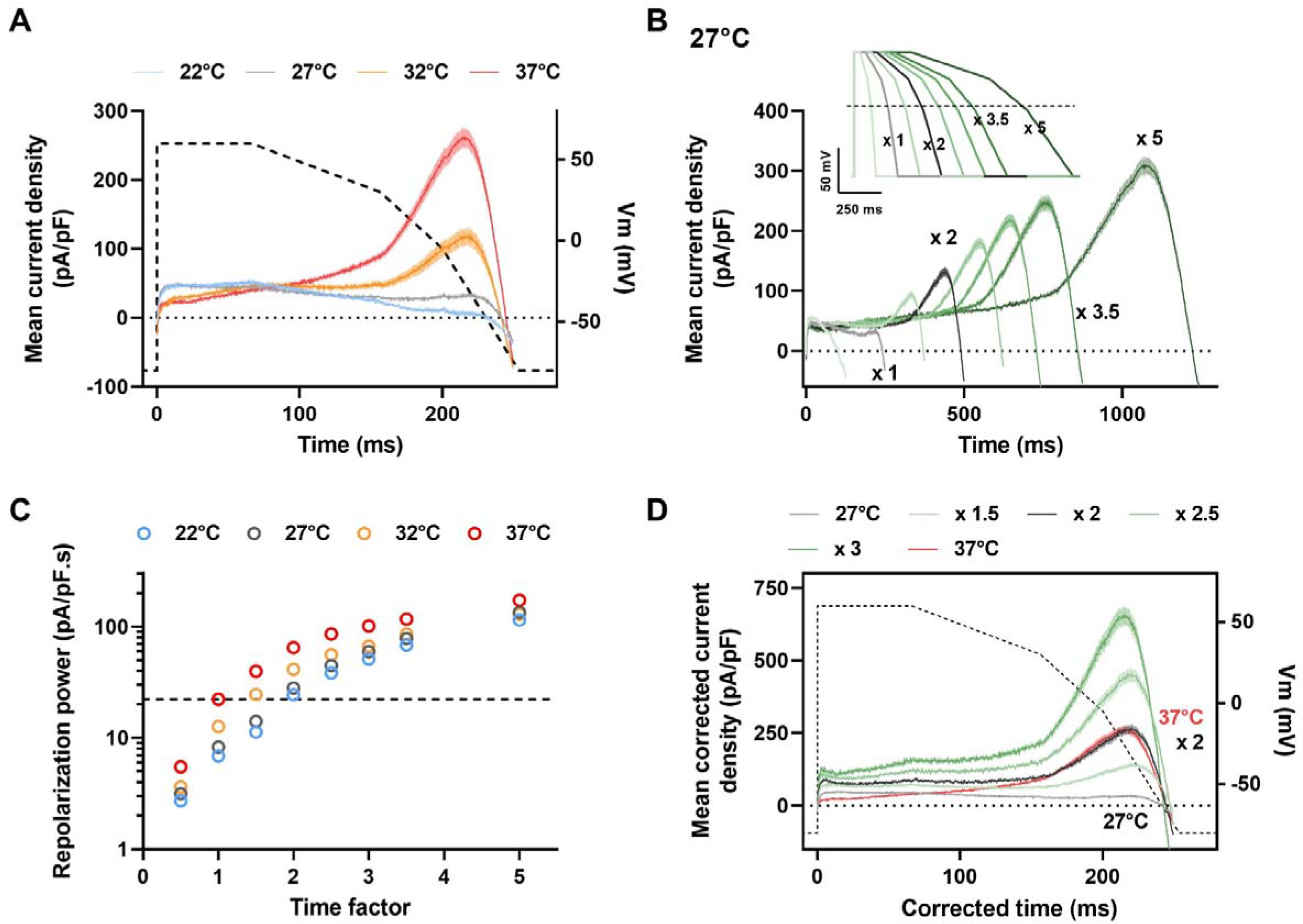
Repolarization power of WT hERG as a function of temperature. A simplified and optimized action potential was applied (AP-clamp) by automated patch clamp system in 384-well plates on HEK293 cells stably expressing hERG. (A) Mean (± SEM) current recordings during AP-clamp at various temperatures (in pA/pF, n = 194, 211, 114, 172 cells at 22 to 37°C, respectively). Dashed line: AP time course (voltage scale: right Y axis). (B) Mean (± SEM) current recordings during AP-clamp at 27°C for AP of various durations (see inset for APs; for currents: n = 168-254 for Time x 0.5 to x 5). The small inward current observed in (A) and (B), when the AP is returning to resting values, is attributed to a contamination of the intracellular solution by the extracellular Tyrode solution, intrinsic to the cell catch process in automated patch clamp (see supplementary information). (C) Mean (± SEM) time integral of the recorded currents densities: repolarization power, at various temperatures *vs.* time factor (n = 155-232, 168-254, 79-133, 144-173 at 22 to 37°C, respectively). Horizontal dashed line: repolarization power at 37°C: 22.3 pA/pF.s. (D) As in (B), at 27°C, after time and current density corrections using various factors from 1.5 to 3 on the respective recordings. For example, for the recordings obtained during the AP of 2 x duration, the time was divided by 2 and the current density multiplied by 2. Note the overlap of the current corrected by the factor of 2 at 27°C (black) with the reference current obtained during the standard AP at 37°C (red).

In order to compensate for reduced hERG channel kinetics at temperatures lower than 37°C, and hence to allow larger current development during AP stimulation, the stimulation protocol was adapted by applying a time factor. For example, when a factor of 2 was used, the AP duration was linearly doubled and the stimulation frequency slowed by 2 (see Figure 2B inset). Time factors varied from 0.5 to 5. Currents were sampled at 5 to 20 kHz. For each set of experiments, results were obtained with the same cell batch, on different plates (one per temperature), on the same day. Only recordings obtained with seal resistance > 500 MΩ, series resistance (R_s_) < 10 MΩ, cell capacitance (C_m_) > 10 pF, and with leak current between 0 and −200 pA at −80 mV before the 4^th^ AP were considered for analysis achieved with an automated R routine. Currents are expressed as current densities in pA/pF. The repolarization power was calculated as the time integral of the absolute current density during the 4^th^ AP.

### Conventional low-throughput electrophysiology

Temperature effects on WT and mutant hERG channels were also investigated on transfected HEK293 cells using conventional patch-clamp. One day after plating, the cells were mounted on the stage of an inverted microscope and constantly perfused by a HEPES-buffered Tyrode solution first maintained at around 20.0°C, at a rate of 3 mL/min. HEPES-buffered Tyrode solution contained (in mmol/L): NaCl 145, KCl 4.0, MgCl_2_ 1.0, CaCl_2_ 1.0, HEPES 5.0, glucose 5.0, pH adjusted to 7.4 with NaOH. Patch pipettes (tip resistance: 2.0 to 2.5 MΩ) were pulled from soda lime glass capillaries (Kimble-Chase). The pipette was filled with an intracellular medium containing (in mmol/L): KCl 100, potassium gluconate 45, MgCl_2_ 1.0, EGTA 5.0, HEPES 10, pH adjusted to 7.2 with KOH. Stimulation and data recording were performed with pClamp 10, an A/D converter (Digidata 1440A) and a Multiclamp 700B amplifier (all Molecular Devices). Currents were acquired in the whole-cell configuration, filtered at 3 kHz and recorded at a sampling rate of 6.9 to 20 kHz. Before membrane capacitance and series resistance 70%-compensation, a series of twenty 30-ms steps to −80 mV was applied from a holding potential (HP) of alternatively −70 mV and −90 mV to subsequently off-line calculate C_m_ and R_s_ values from the recorded currents. A 3-step protocol was used to test the current rundown/runup (HP =-80 mV, 1^st^ pre-pulse: +40 mV during 1 s, 2^nd^ pre-pulse: −100 mV during 15 ms, test-pulse: +40 mV during 500 ms, every 5 s). Then, an AP was used to clamp the voltage, that derived from the same O’Hara and Rudy model^31^ as in automated patch-clamp. Unlike for automated patch-clamp, intracellular fluoride or high external Ca^2+^ concentration is not required to easily obtain Gigaseals. Therefore, the AP stimulation was corrected by the calculated liquid junctional potential only (see Figure 3A, the dashed line representing the AP stimulation)^24^. Three repeated APs were enough to stabilize the resulting I_hERG_ current time course. From around 20.0°C, Tyrode temperature was gradually raised until it reached 32.0°C in cell bath and AP-clamp recording was performed every 2.0°C. As in high-throughput experiments, in order to compensate for reduced hERG channel kinetics at lower temperatures, the stimulation protocol was adapted by applying a time factor, from 1 to 3: the same AP stimulation file (Axon text file: *.atf, compatible with Clampex 10 of the pClamp software suite, Axon Instruments, CA, USA) was used at various frequencies. We kept the typical quality control parameters of conventional patch-clamp to select the cells: seal resistance > 1GΩ, series resistance < 10 MΩ at the beginning and the end of the recording, and with less than 200 pA of leak current at −80 mV before the 3^rd^ AP that was used to calculate the repolarization power. At the end of the experiment, the pipette potential offset was checked and recordings presenting offset > 5 mV were discarded. Currents are expressed as current densities in pA/pF. It has to be mentioned that only cells presenting detectable hERG currents were analyzed in order to be able to compare repolarization powers at various temperatures in the same cell. Data were analyzed and compiled using ClampFit 10 of the pClamp software suite, Microsoft Excel, and Prism (GraphPad Software, CA, USA). Statistical analyses were processed using Prism with Wilcoxon matched-pair signed rank test, or Mann-Whitney test, when appropriate. Significance level was set to 0.05.

**Figure 3:**
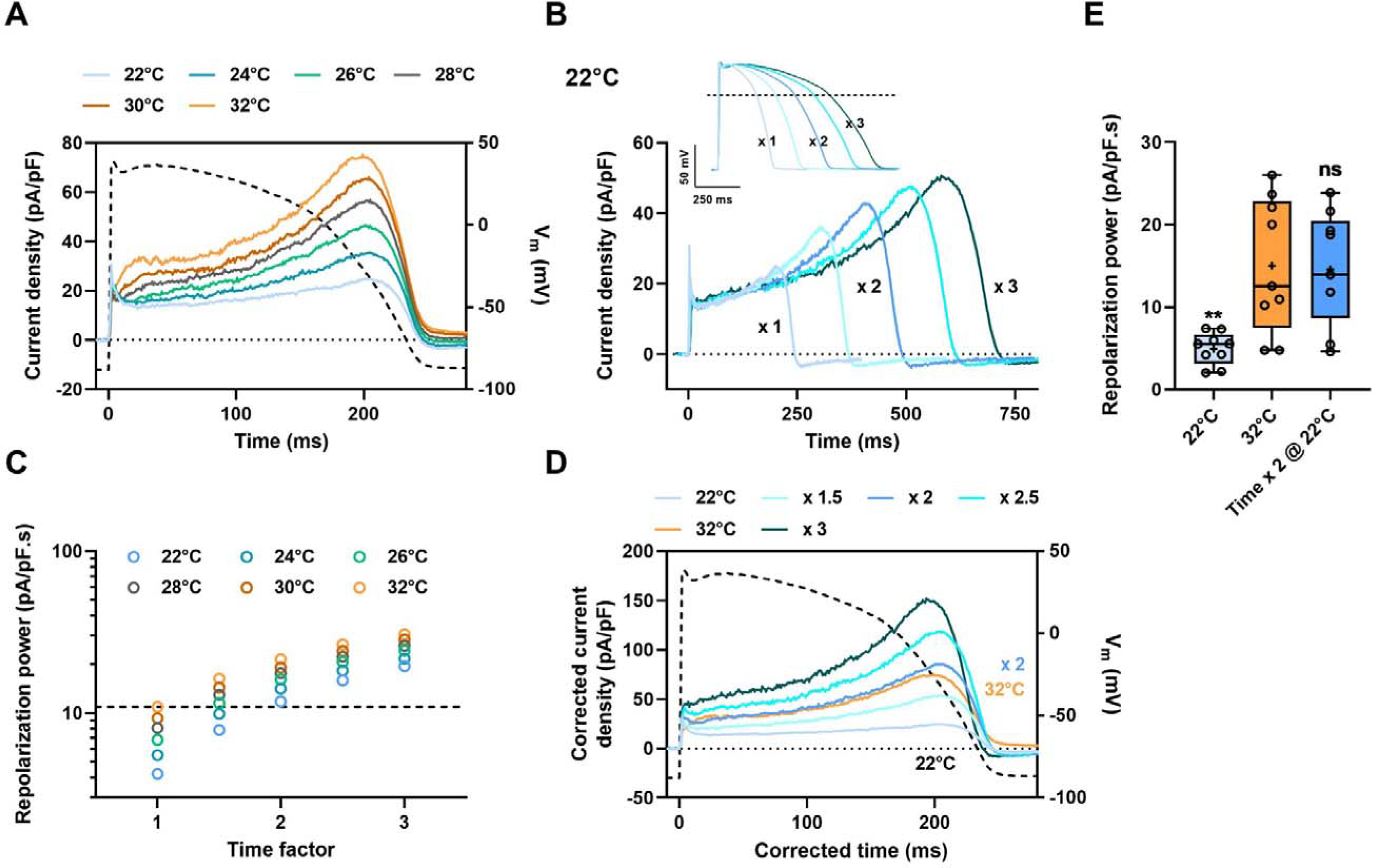
Repolarization power of WT hERG as a function of temperature in HEK293 cells transiently expressing hERG. A model action potential was applied (AP-clamp) using conventional patch-clamp. (A) Typical current recordings on a single cell during AP-clamp at various temperatures. Dashed line: time course of the model AP^31^ (voltage scale: right Y axis). (B) Current recordings during AP-clamp at 22°C (same cell as A) for AP of various durations (inset). (C) Corresponding repolarization power at various temperatures *vs.* time factor. Horizontal dashed line: reference repolarization power at 32°C: 10.9 pA/pF.s for this cell. (D) As in B, at 22°C, after time and current density corrections using various factors from 1.5 to 3 on the respective recordings. For the recordings obtained during the AP of x 2 duration, the time has been divided by 2 and the current density multiplied by 2. Again, note the overlap of the current at 22°C, corrected by 2, as in figure 1D (blue) with the reference current obtained during the standard AP at 32°C (orange). (E) Tukey plot of the repolarization powers recorded at 32 and 22°C during the standard AP or AP of doubled duration (n = 9). *vs.* 32°C: ns: non-significant; **: *P* < 0.01.

## Results

### Temperature dramatically affects the success rate in high-throughput automated patch-clamp system

To illustrate the impact of temperature on the success rate of hERG recordings, we first investigated the seal quality as a function of temperature using stable cell lines expressing hERG channels. For that purpose, we used the automated patch-clamp system that provides robust information on the temperature effect thanks to the great numbers of cells that can be recorded. As illustrated in Figure 1A, the percentage of HEK293 cells with high-quality seal resistance ≥ 1 GΩ decreased significantly when the recording temperature is increased. The success rate dropped from 38 to 12% when measured at 27 and 37°C, respectively. In this case, one could consider that the success rate at 37°C is acceptable although very low. However, if one aims to characterize hundreds of hERG variants, the most realistic approach is to use transient expression which may add another fragilizing factor to seal quality. To test this, we electroporated HEK293 cells with a plasmid over-expressing hERG WT or mutant and estimated again the effect of temperature on seal quality. The percentage of cells reaching a seal resistance ≥ 1 GΩ at 27°C reached 18% but only 21 cells (5.4% of the plate wells) exhibited hERG currents above 50 pA. In contrast, at 37°C, the percentage of cells with ≥ 1 GΩ seal resistances dropped to 5% and none of them displayed a hERG current. Including cells with seal resistance < 1 GΩ but ≥ 500 MΩ improved marginally only the rate of cells presenting measurable K^+^ currents (3 cells at 37°C - Figure 1B). On the other hand, for HEK293 cells stably expressing hERG channels, the success rate of 38% at 27°C increased to 77% when the seal resistance limit dropped from 1 GΩ to 500 MΩ. Of note, this seal quality remains largely above previously accepted ones by others during automated patch-clamp experiments (≥ 200 MΩ)^29^.

**Figure 1:**
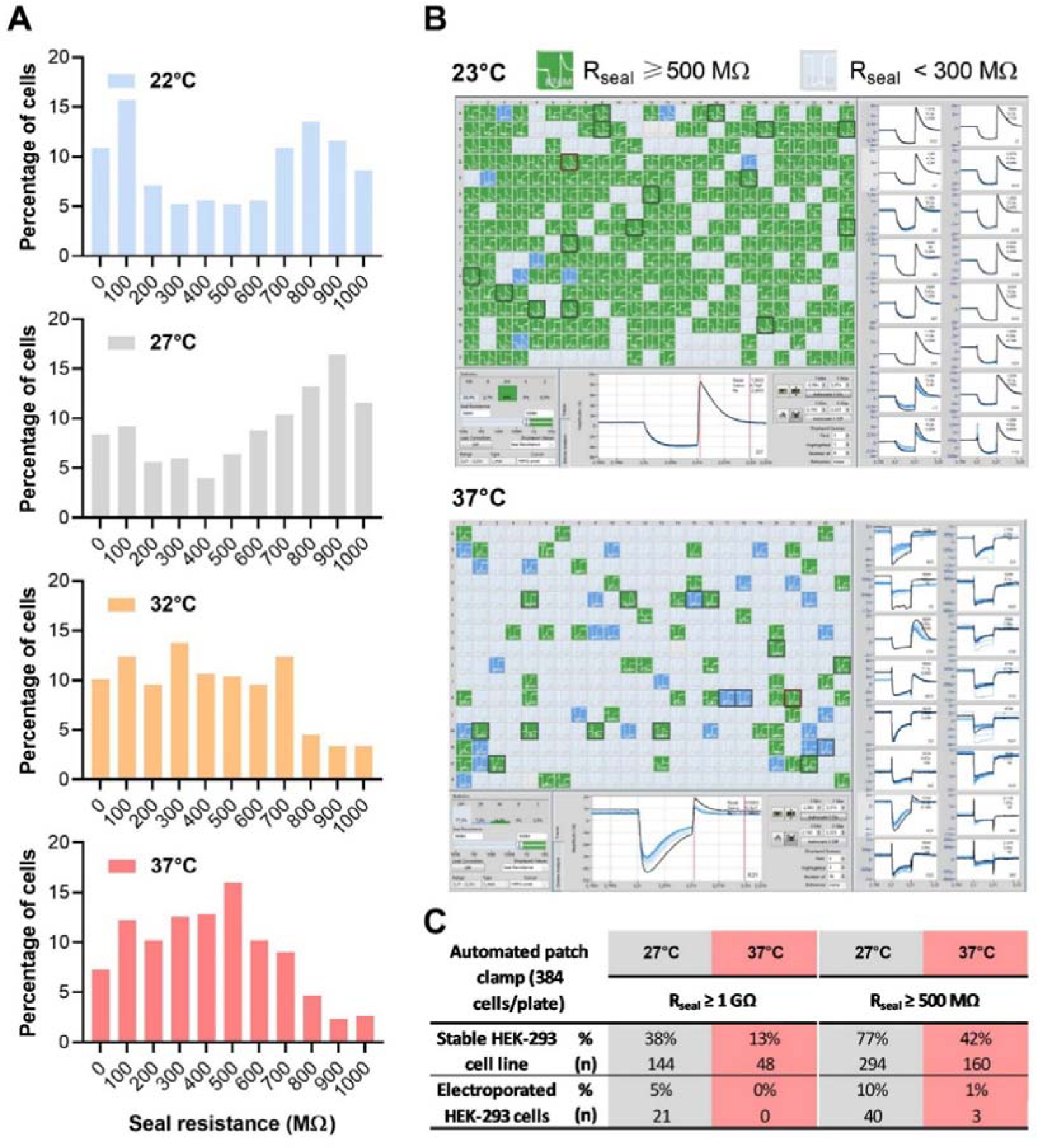
Evolution of the seal resistance (R_seal_) with increasing temperatures. (A) R_seal_ value distribution for HEK293 cells stably expressing WT hERG channels (one 384-well plate per temperature) using automated patch-clamp system. (B) Graphical user interface illustrating single-cell recordings from transiently transfected HEK293 cells at 23 and 37°C, 30 h post-transfection (electroporated cells, one 384-well plate per temperature). Each recording well is visualized and color-coded based on user-defined quality criteria. A single-cell recording is considered as successful when R_seal_ ≥ 500 MΩ (green). (C) Summary of percentage of cell with measurable hERG current in various conditions. The dramatic drop of recordings with R_seal_ ≥ 500 MΩ at 37°C precludes the hERG current measurements at physiological temperature on transiently transfected cells using automated patch-clamp system

However, even the medium-quality experimental conditions with seal resistance values above 500 MΩ are not sufficient to reasonably increase data production yield at physiological temperature if one desires to tackle the biophysical properties of hundreds of hERG variants. Thus, an alternative protocol is needed to predict at room temperature the *in vivo* impact of variants. As validation, we compared currents obtained at room and physiological temperatures using automated patch-clamp system for HEK293 cells stably expressing hERG channels and conventional patch-clamp technique for cells transiently over-expressing hERG variants, choosing, in this latter case, gold-standard quality criteria to accurately evaluate the temperature effects.

### The repolarization power of WT hERG channels depends on the temperature

As mentioned above, the hERG current recorded during an AP-clamp protocol can be used to estimate, by mathematical integration, the total amount of K^+^ ions crossing the membrane during the AP. We named this value the repolarization power, since it represents the hERG contribution to the late repolarization phase of the AP. hERG being a voltage-gated channel, I_Kr_ steady-state is never reached during the cardiac AP. In the cardiomyocyte, its amplitude and contribution to repolarization depends on contribution of the other depolarizing and repolarizing currents, all of them shaping the AP time course and in return, all of them being shaped by the transmembrane voltage change. To standardize any estimation of hERG repolarization power during the action potential, it is necessary to clamp most of the factors regulating hERG contribution to the AP. This is achieved *in vitro* with a standard AP stimulation. Then, the repolarization power is a good representation of the channel activity because it takes into account, altogether, the number of channels expressed at the plasma membrane and the voltage-dependence and kinetics of hERG current. Thus, this global index allows reporting biophysical changes introduced by a given condition or hERG channel mutation.

Standard AP shape has been logically defined at 37°C and the resulting I_hERG_ current recorded at 37°C is the reference for the I_Kr_ time course during the cardiac action potential *in vivo*. However, for the sake of efficiency we had to use lower temperatures. Decreasing the recording temperature affects the biophysical processes at work within the channel and thus should affect the repolarization power. We took advantage of the automated patch-clamp system to efficiently characterize the effect of temperature on AP-evoked I_hERG_ and repolarization power in HEK293 cells stably expressing WT hERG. Figure 2 illustrates the results obtained at various temperatures for the WT hERG channels, in a first set of experiments. As shown, the amount of K^+^ ions crossing the cell membrane during an AP indeed decreased with decreasing temperatures (Figure 2A). At 27°C, a temperature compatible with high recording yields, the hERG contribution to AP late repolarization dropped. Since seal quality and high temperature are mutually exclusive, we had to envision a strategy to restore at 27°C a repolarization power equivalent to the one observed at 37°C during the standard AP.

### The reference repolarization power value of hERG channels can be restored by extending the AP duration

In order to obtain the reference repolarization power of hERG channels at room temperature, we need to compensate the effects of temperature on hERG kinetics. To do so, we arbitrarily ’shrunk’ or ’dilated’ the time with a given factor on the AP protocol duration (Figure 2B, inset). The effect of this time factor on the AP was studied at a constant temperature of 27°C. As shown in Figure 2B, I_hERG_ has more time to develop with longer APs. As a result, the average repolarization power increased and the same effects were observed at each tested temperature (Figure 2C). For the sake of comparison, we included in this representation, reference I_hERG_ and reference repolarization power (time factor x1) (Figure 2A, red trace, and Figure 2C, dashed line, respectively). At 27°C, the repolarization power was logically lower than at 37°C when time factor was 1 (Figure 2C). With a time factor of 2, the repolarization power at 27°C approximately matches the one obtained at 37°C. Perfect matching was inferred by fitting the data at 27°C in Figure 2C giving an exact value of 1.78. When we repeated the experiments at 27°C and 37°C in a second set of experiments, similar results were observed (time factor of 2.18, Supplemental figure 3).

When Vandenberg *et al.* investigated the hERG temperature sensitivity, they calculated a temperature coefficient (Q_10_, factor expressing the change observed when the temperature is increased by 10°C) of 1.4 for the whole cell conductance and of 1.7 to 2.6 for activation, deactivation, inactivation and recovery from inactivation, the lasts being the largest^26^. Here we postulated a unique value of 2 for every parameter including whole cell conductance. When the time factor was used to correct both kinetics and current amplitude *i.e.* as a unique ’quasi-Q_10_’ of all channel parameters, the time course of the corrected current recorded at 27°C with a time factor of 2 was the most similar to that at 37°C (Figure 2D and Supplemental figure 3D). In summary, the dilatation of the time factor had no consequences on the shape of the repolarizing current since, in these plots, the two currents display the exact same kinetics. This result further validates this approach of AP duration variation for the accurate prediction of the repolarization power of hERG channels.

In conclusion, slowing the AP protocol by a factor of 2 generates hERG current at 27°C and its corresponding repolarization power that predict the ones obtained at 37°C.

**Supplemental figure 3:**
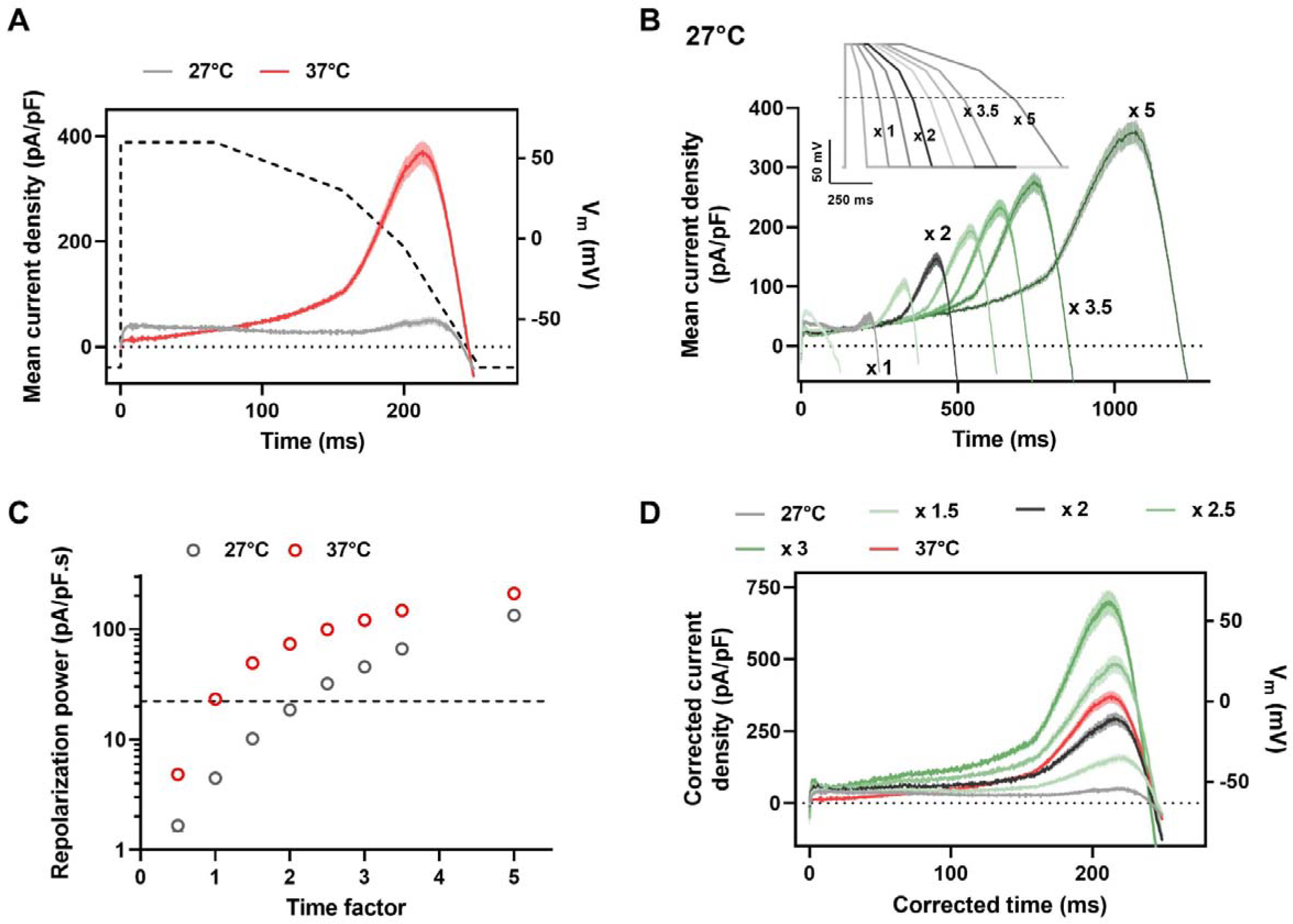
Repolarization power of WT hERG as a function of temperature – Replicate. A simplified action potential was applied (AP-clamp) on HEK293 cells stably expressing WT hERG by automated patch clamp system in 384-well plates. (A) Mean (± SEM) current recordings during AP-clamp at 27 and 37°C (in pA/pF, n = 123 and 150 cells, respectively). Dashed line: time course of the simplified AP (voltage scale: right Y axis). (B) Mean (± SEM) current recordings during AP-clamp at 27°C for AP of various durations (see inset for APs; n = 59-230 for Time x 0.5 to x 5). (C) Mean (± SEM) repolarization power at 27 and 37°C *vs.* time factor (n = 59-230 and 107-158, respectively). Horizontal dashed line: repolarization power at 37°C: 23.2 pA/pF.s. (D) As in (B), at 27°C, after time and current density corrections using various factors from 1.5 to 3 on the respective recordings. For the recordings obtained during the AP of x 2 duration, the time was divided by 2 and the current density multiplied by 2. Note the overlap of the current corrected by the factor of 2 at 27°C (black) with the reference current obtained during the standard AP at 37°C (red).

### A two-fold extension of the AP duration is also valid for two hERG variants

Then, we challenged the determined value in the case of 2 variants of hERG linked either to aLQTS (p.R328C) or to short QT syndrome (SQTS, p.D591H) that we formerly characterized^22, 24^. For that purpose, we used the conventional patch-clamp technique on hERG transiently over-expressed in HEK293 cells. As for automated patch-clamp, we frequently failed to keep the Giga-seal until the temperature reached 37°C. Therefore, we investigated the factor validity between 22 and 32°C. Figure 3 illustrates first the results obtained on WT hERG in a typical cell. Again, the current increased with temperature or AP duration (Figure 3A and B, respectively). Also, the current density profiles were similar when recorded at 32 °C and corrected at 22°C with a factor of 2 (Figure 3C and D). On average, we calculated a factor of 1.98 ± 0.06 (n = 9) between currents recorded at 22 and 32°C. The repolarization powers extracted from data at 32°C, and 22°C during a time-doubled AP, were not significantly different (15.1 ± 2.7 pA/pF.s and 14.6 ± 2.3 pA/pF.s, respectively, Figure 3E). By comparison, when the current was recorded at 22°C with the original AP (Time factor = 1), the calculated repolarization power value was much smaller than that obtained with the same AP at 32°C (5.0 ± 0.7 pA/pF.s), as observed with the automated patch clamp.

When applied on cells overexpressing the p.R328C hERG variant, the current density profiles of p.R328C hERG current time course recorded at 32 °C could be replicated at 22°C after correction with a time x 2 modified AP (Figure 4B), as for WT hERG channels. Profiles were not fully matching in the case of p.D591H hERG variant. At 22°C the I_hERG_ peak was shifted toward the end of the AP (Figure 4E). Despite this shift, the effect of p.D591H observed at 32°C could be still visualized at 22°C, when p.D591H hERG normalized corrected current time course was compared to that of WT hERG (Supplemental figure 4C). Again, values of 1.92 ± 0.12 (n = 6) and 1.92 ± 0.06 (n = 8) were calculated (Figure 4A and 4D for typical cells). As for WT, the repolarization power extracted from data at 32°C, and 22°C during a time-doubled AP, were not significantly different for each variant (Figure 4C and 4F, n = 6 and 8, respectively).

**Figure 4:**
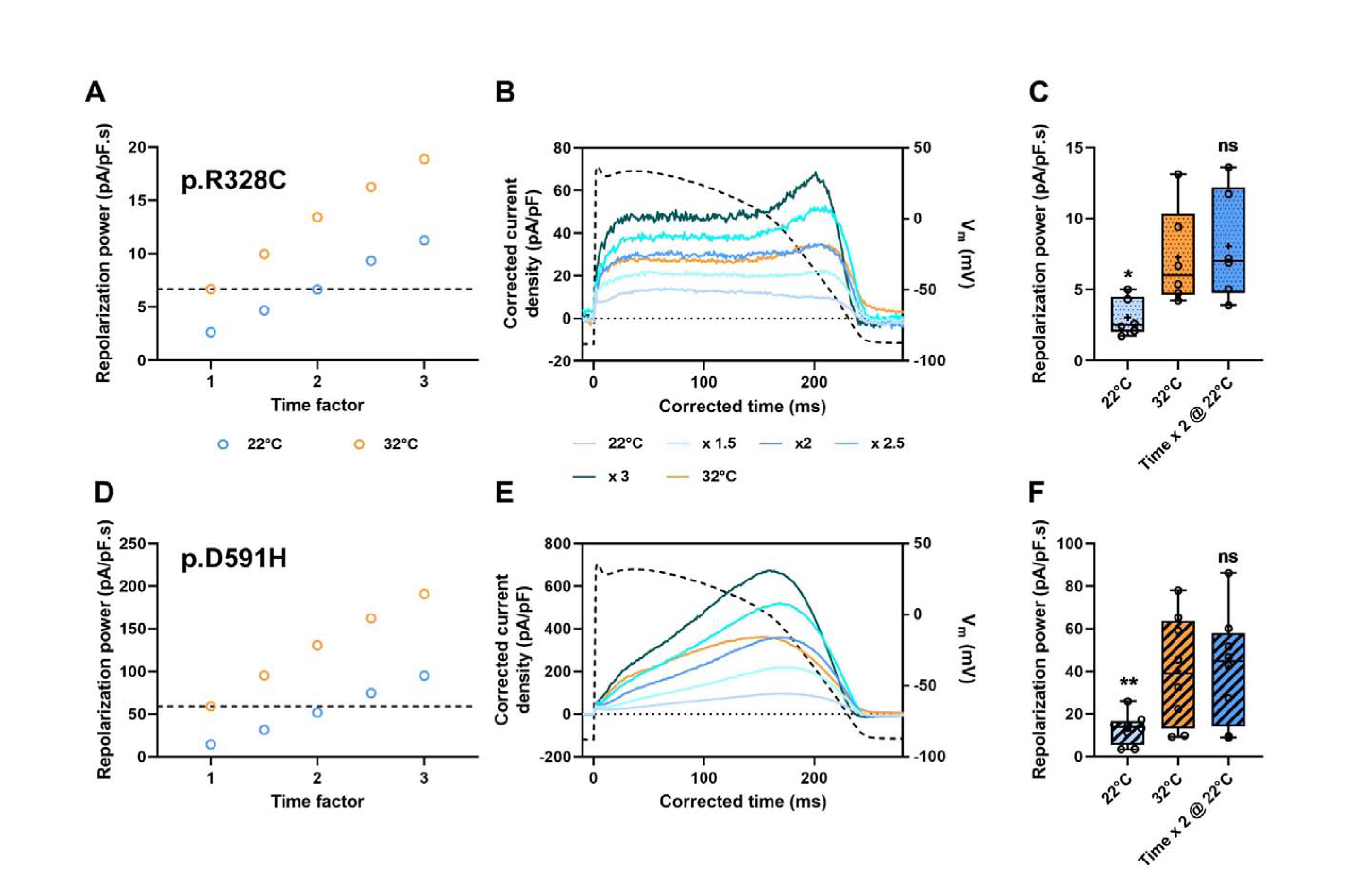
Repolarization power of LQTS (A) and SQTS (B) hERG as a function of temperature. Action potential clamp was applied (AP-clamp) on HEK293 cells transiently expressing p.R328C (A-C) or p.D591H (D-F) hERG variant. (A & D) Repolarization power at various temperatures *vs.* time factor in typical cells. Horizontal dashed line: reference repolarization powers at 32°C: 6.7 and 59.1 pA/pF.s for p.R328C and p.D591H hERG, respectively. (B & E) Current recordings during AP-clamp at 22°C (blue; same cells as A & D, respectively) for AP of various durations after time and current density corrections using various factors from 1.5 to 3 on the respective recordings; reference current recording during AP-clamp at 32°C (orange). (C & F) Tukey plot of the repolarization powers recorded at 32 and 22°C during the standard or AP of doubled duration, for each variant (n = 6 and 8 cells for p.R328C and p.D591H, respectively). *vs.* 32°C: ns: non-significant; *: *P* < 0.05; **: *P* < 0.01.

The aim of this experimental set was not to characterize the effects of variants on current time course or repolarization power compared to WT. Since only cells with detectable hERG currents were included in the analysis, such selection tends to blunt the potential effect of a mutation. However, it is worth noting that despite this cell selection, effect of p.R328C mutation, when compared to WT, could be clearly detected on the I_hERG_ repolarization power value (Supplemental figure 4B) determined at 22°C using a time x 2 modified AP but not using an AP of normal duration at the same temperature.

Altogether, these data indicate that a time factor of 2 may be used to efficiently determine the repolarization power at 22°C, highly similar to that developed at 32°C during an AP-like stimulation. This time factor should be valid to predict WT and variant hERG repolarization power at physiological temperature.

**Supplemental figure 4:**
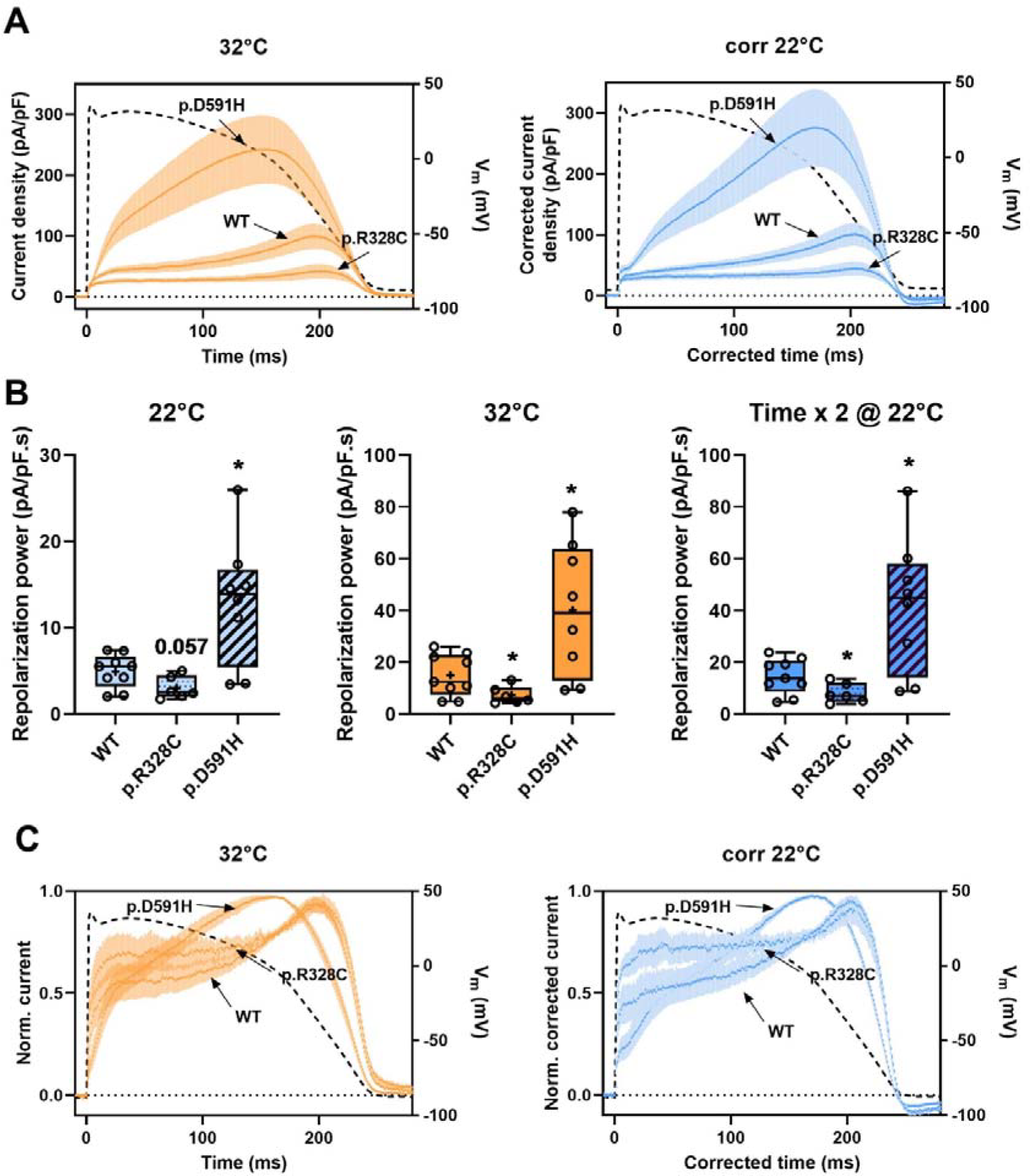
Repolarization power of WT, LQTS (p.R328C) and SQTS (p.D591H) hERG as a function of temperature – Comparison. AP clamp was applied on HEK293 cells transiently expressing WT or hERG variants (same cells as  Figures 3 and 4). (A) Mean (± SEM) current recordings during AP-clamp at 32°C (orange; n = 9, 6 and 8 for WT, p.R328C and p.D591H, respectively) and for AP of 2x duration at 22°C after time and current density corrections using the coefficient of 2 on the respective recordings (blue). (B) Tukey plots of the repolarization powers recorded at 32 and 22°C during standard or AP of doubled duration. *vs*. WT: *: *P* < 0.05 or *P* value when non-significant. (C) As in (A) after normalization to the maximum current density value.

## Discussion

Comparing the amount of potassium ions exiting cells expressing WT and a variant hERG channel during an action potential can be used as an index of the variant pathogenicity. However, such comparison needs to be operated at physiological temperature in order to faithfully simulate the channel behavior, such temperature being far from optimal in high throughput patch-clamp. In an attempt to get a useful and simple repolarization power index of *KCNH2* genetic variants compatible with high throughput phenotyping pipeline and as informative as that obtained at physiological temperature, we found that using a time factor of 2 on the AP duration used as voltage stimulation was needed to compensate a 10°C decrease of the experimental temperature.

Since the profile of the AP-clamp generated I_hERG_ strongly depends on the open state occupancy of the channel, the repolarization power encapsulates all the kinetics parameters in a synthetic index. Despite the diversity of the temperature effects on these different rates^26, 28^, we propose that a single factor can sufficiently compensate for the effects of temperature change and allows to extrapolating the repolarization power value at physiological temperature from the value obtained at lower temperature. Alternatively, the method established by Lei *et al.*^28^ has demonstrated its efficacy to characterize the 4 different kinetic rates for hERG channels at different potentials, corresponding to activation, deactivation, inactivation and recovery rates, consequently to infer the parameters describing the voltage dependence of steady states and time constants of the activation and inactivation gates, and eventually, to predict the effects of temperature^27, 28^. But this approach is based on a Hodgkin-Huxley model that may oversimplify the actual channel behavior. The hysteretic behavior of the activation observed, for example, by Jones^36^ cannot be simulated using Lei’s model. In addition, if the Q_10_ formulation is chosen, Lei and collaborators calculated 4 different Q_10_ values for the 4 kinetic rates^28^. In fact, they deducted a higher degree of complexity, showing that the temperature dependence had also a voltage-dependent component.

We are aware that using a single correcting value to summarize the effects of temperature on hERG current is an approximation^25, 26, 28^. However, it appears to be reliable in the strictly limited context of the AP course. In our study, the repolarization power index that we propose is unbiased *i.e.* model-free, and is directly related to the electrophysiological conditions of the hERG activity in the heart: during the action potential. Using a unique time factor of 2 on the AP stimulation at room temperature as an approximation, we obtained satisfactory results to simulate the current profile during the AP at more physiological temperatures (+10°C). The aim of our work differs from that of Lei, in the sense that we only want to identify the net effects of a given variant on the repolarization reserve of the cardiomyocytes and not to their full and detailed description.

In the case of two different mutants, applying the same time factor at low temperature was sufficient to obtain the same repolarization power value as that at higher temperature. This procedure may thus become compatible with the use of an automated patch-clamp system for the determination of the repolarization power by simply shifting to lower values the range of temperatures one need to work with. In some cases, though, the temperature dependence may be impaired by a mutation and this kind of information may be missed at room temperature. The p.A558P and p.F640V hERG missense mutations have been reported in patients suffering from fever-induced polymorphic ventricular tachycardia^37^. Investigating the trafficking and function of the hERG channels in the heterozygous condition showed a limited trafficking associated with a largely impaired K^+^ current due to the dominant-negative effect of these mutations. In addition, the current generated by co-expression of mutant plus WT hERG channels did not increase to the same extent as the WT current at higher temperatures (fever) in transfected HEK293 cells, exacerbating the effects of these mutations. This limit would be a problem when the variant activity is preserved at room temperature and impaired at higher temperature, but this case may be very rare since, up to now, no such a variant has been reported. However, whatever the method used to extrapolate the effects of temperature, the information related to the temperature sensitivity changes induced by a variant will be lost when evaluated at low temperature. Actually, the effects of p.A558P and p.F640V variants were detected already at room temperature. The following question is: do we miss this information to evaluate the molecular pathogenicity of a given variant? In fact, getting the repolarization power index at low temperature would underestimate it, only.

A much more common limitation is revealed by the study of p.D591H variant. Here, we show that the early contribution of the current during the AP is blunted at lower temperature. This is most probably the case with many loss-of-function mutations acting on activation and/or inactivation gating. It may be of importance in situations when the repolarization power is not severely impaired when compared to WT. Nonetheless, the superimposition of the corrected current time course of WT and p.D591H hERG channels at 22°C after normalization on maximal amplitude still unmasks these effects (Supplemental figure 4C). Therefore, it may be of interest to find an additional way to automatically detect this behavior. To do so, a second index could report the differences to WT mean current during the AP time course. Anyhow, it would require further validations by studying a larger number of variants, though.

Finally, we purposely chose the p.R328C as the aLQTS variant since divergent results were obtained in three different studies, two of them originated from our laboratory^22–24^. Investigations when overexpressed in HEK293 or CHO cells failed to detect any current changes at 22°C, whereas we observed a reduction of above 60% in COS-7 cells at 37°C^22^. Here, evaluating the effects of temperature on the repolarization power, we observed that the significance of the decrease when compared to the WT index could be reached at 32°C and at 22°C with the time-adapted AP but not with the standard AP at 22°C (Supplemental figure 4B). We have to keep in mind though that the purpose of the study was not to validate the index robustness in this last condition (standard AP duration at 22°C). However, using larger samples of WT and p.R328C hERG transfected cells, including cells without detectable hERG currents, in our previous evaluation of 13 LQT2 variants, we observed a significant repolarization power decrease, using the normal AP at 22°C^24^. Therefore, it appears that despite the partly blunted effects at 22°C, the pathogenicity of such a variant can be unmasked by the new protocol tested here.

In summary, we present a simple, reliable and practical protocol to measure the reference repolarization power of a given hERG mutant at room temperature. We propose the reference repolarization power, relative to the WT one, as a functional index of variant pathogenicity, in the range of the cardiac action potential, more informative than the relative current density suggested by Jiang *et al*.^38^.

